# Folding of VemP into translation-arresting secondary structure is driven by the ribosome exit tunnel

**DOI:** 10.1101/2021.04.15.440051

**Authors:** Michal H. Kolář, Gabor Nagy, John Kunkel, Sara M. Vaiana, Lars V. Bock, Helmut Grubmüller

## Abstract

The ribosome is a fundamental biomolecular complex responsible for protein production in cells. Nascent proteins emerge from the ribosome through a tunnel, where they may interact with the tunnel walls or small molecules such as antibiotics. These interactions can cause translational arrest with notable physiologic consequences. Here, we studied the arrest caused by the regulatory peptide VemP, which is known to form an *α*-helix in the ribosome tunnel near the peptidyl transferase center under specific conditions. We used all-atom molecular dynamics simulations of the entire ribosome and circular dichroism spectroscopy to study the driving forces of helix formation and how VemP causes the translational arrest. To that aim, we compared VemP dynamics in the ribosome tunnel with its dynamics in solution. We show that the VemP sequence has a low helical propensity in water and that the propensity is higher in more hydrophobic solvents. We propose that helix formation within the ribosome is driven by the tunnel environment and that a portion of VemP acts as an anchor. This anchor might slow down VemP progression through the tunnel enabling the *α*-helix formation, which causes the elongation arrest.

## Introduction

In all organisms, the proteins are synthesized by ribosomes. Each ribosome consists of several strands of ribonucleic acid (RNA) and proteins organized in two ribosomal subunits. Ribosomes translate the genetic information stored in the messenger RNA into a sequence of amino acids (AAs). The small subunit reads the information, the large subunit catalyzes peptide bond formation. One by one, AAs are delivered to the ribosome by transfer RNAs (tRNAs) and are covalently attached to the nascent peptide chain (NC) at the peptidyl-transferase center (PTC) of the ribosome.

The PTC is buried deep within the large subunit and each NC leaves the ribosome through a 10-nm long exit tunnel. Recent studies have suggested an active role of the tunnel in protein synthesis and translation regulation. ^1,2^ Some NCs interact with the tunnel walls such that peptide-bond formation is slowed down or inhibited completely. This elongation arrest can have many physiological consequences. For instance, some stalling NCs were shown to stall on faulty mRNAs^3,4^ or to serve as chemical^5–8^ or mechanical^9,10^ sensors. In addition, a large class of antibiotics acts by binding inside the ribosome tunnel and causing ribosome stalling. ^11^

The inherent flexibility of NCs complicates their structure determination by classic biophysical techniques such as X-ray crystallography or cryogenic electron microscopy (cryo-EM). Nevertheless, several ribosome nascent-chain (RNC) complexes have been resolved, taking advantage of the elongation arrest.^5–7,12–16^ Computer simulations can be used to complement structural data by dynamics and energetics and to probe the unstalled conformations. ^15,17–19^

A recently discovered stalling NC is *Vibrio* export-monitoring peptide (VemP). ^10^ In the marine-estuarine bacterium *Vibrio alginolyticus*, VemP regulates the expression of a gene variant via an elongation arrest, thus enabling the bacterium to survive in fresh as well as in sea water.^10^ The structure of the nascent VemP peptide in the tunnel is controlled by an external mechanical force acting on the N-terminus of the peptide. When the force, normally generated by the translocon channel, is acting on VemP at the exit of the tunnel, VemP is most likely extended in the ribosome tunnel and translation proceeds undisturbed.^19,20^ In the absence of the force, however, VemP adopts a compact and well-defined secondary structure, as discovered by cryo-EM. ^16^ The cryo-EM atomic model of the ribosome-VemP complex, determined at an average resolution of 2.9 °A, revealed that VemP folds into two *α*-helices connected via an S-shaped loop (Fig. 1a). The inner helix is located at the PTC, whereas the outer helix occupies a wider corridor located beyond the constriction of the tunnel formed by extensions of ribosomal proteins uL4 and uL22 (Fig. 1b).

**Figure 1:**
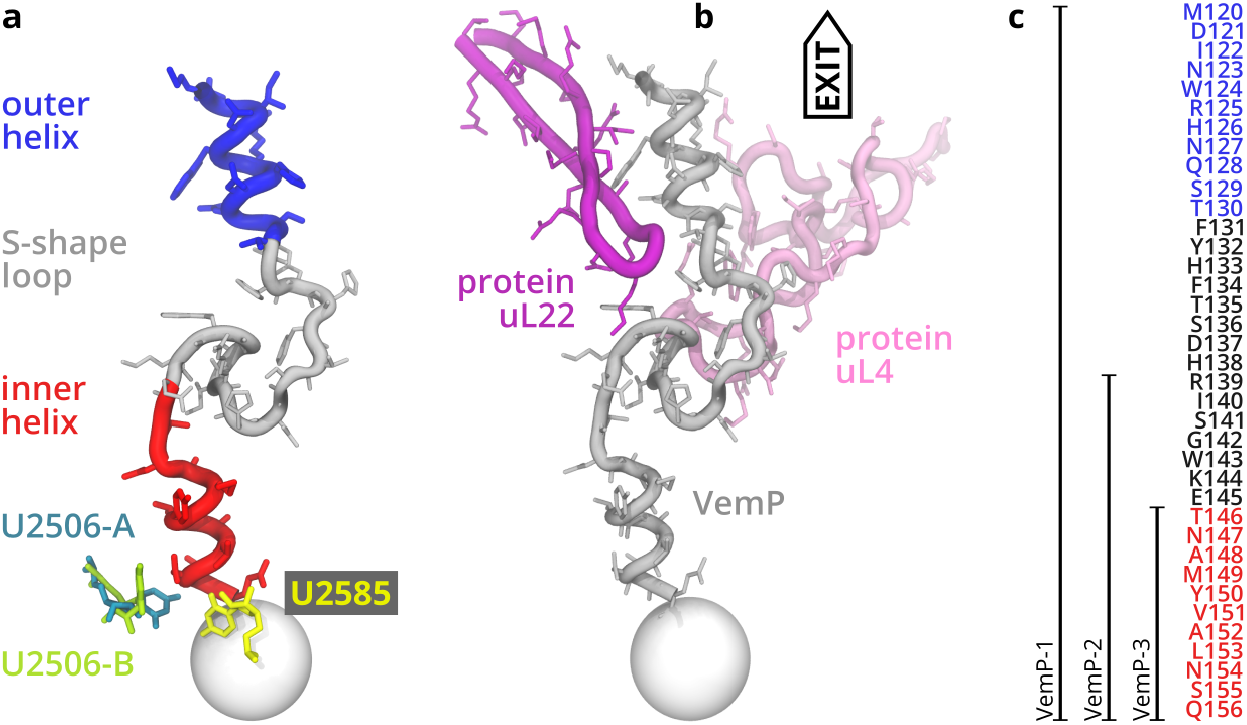
VemP overview. **a)** Relevant VemP parts with the critical nucleotides at the peptidyl trasferase center (sphere). Two substates of U2506 are denoted -A and -B. **b)** VemP stalling conformation ^16^ in the context of two exit-tunnel proteins uL4 and uL22 **c)** Primary structure and a schematic representation of the constructs studied.

Based on the cryo-EM data, it was proposed that the inner helix induces a conformational change of the PTC nucleotides U2506 and U2585 (*E. coli* numbering). This conformation is sterically incompatible with the accommodation of tRNA into the A site and thus prevents the delivery of the new amino acid^16^ (phenylalanine in the VemP case). Moreover in the cryo-EM model, U2506 was observed in two substates, denoted *A* and *B*, each with an occupancy of 50%. The nucleotide U2506 in the A substate occupies the volume of the incoming Phe, and in the B substate U2506 is situated on the side of the VemP inner helix. The role of the two substates is not fully understood, however.

VemP contains 159 amino acids and stalls when the codon of Q156 resides in the P site. Radiolabeled toeprinting *in vitro* experiments of single-point VemP mutants ^10^ and *in vivo* LacZ assay^21^ of single-point mutations showed remarkable VemP sensitivity to mutations. The studies found a periodic appearance of the functionally important AAs, which agrees with the helical structure of the stalling motif. The stalling motif, defined by about 20 AAs (136–156), is by more than 10 AAs larger than other stalling peptides like SecM,^22,23^ TnaC,^5,6^ ErmBL, ^15^ or MifM.^9^

Here, we address the question of which molecular driving forces cause VemP *α*-helix formation in the ribosome tunnel. Previously, it was argued that the tunnel features “folding zones”, where a compaction of NC occurs. ^24^ The region nearest to the PTC was identified as having the strongest compaction effect. Also, a coarse-grained off-lattice model in a non-specific confined space suggested that a helix formation is favorable, and entropically driven. ^25^ This is, however, in contradiction with computer simulations of Sorin and Pande,^26^ who argued that helices are destabilized by cylindrical (carbon nanotube-like) space. The dominant role in the confinement was attributed to the entropy of water.^26^

Here, we investigated the dynamics of VemP in the ribosome as well as outside, in solution. To determine the inherent secondary structure (SS) propensities of VemP, we performed all-atom molecular dynamics (MD) simulations of several constructs of the wild-type VemP complemented with circular dichroism (CD) spectroscopy measurements in various solvents. In the ribosome, we finally investigated the role of the U2506 substates in the elongation arrest by running separate sets of MD simulations in both substates.

## Methods

### Studied Molecular Systems

We studied three VemP constructs whose sequences are denoted by VemP-1, VemP-2, and VemP-3 (Fig. 1c). VemP-1 covers residues M120–Q156, which were resolved in the ribosomal tunnel by cryo-EM,^16^ and comprise the inner helix, S-shape loop and the outer helix. A shorter VemP-2 covers residue S141–Q156, which span the inner helix and a part of the loop. The shortest VemP-3 spans only the inner helix and contains residues T146–Q156.

The initial structure of the ribosome-VemP complex was taken from the Protein Data Bank (PDB 5NWY).^16^ It contains an *E. coli* ribosome, the P-site tRNA^*Gln*^ with a part of the wild-type VemP nascent chain (residues 120–156). Because all nucleotides are modeled as the canonical nucleotides (A, C, G, U) in the cryo-EM structure, we added the previously determined nucleotide modifications to the ribosomal RNA using the structure by Fischer et al. (PDB 5AFI)^27^ as a template. In the structure of the ribosome-VemP complex, no A-site tRNA was present. The trajectories of this complex obtained by MD simulations are denoted by *VemP-1R*.

From this initial RNC complex, we prepared an alternative structure containing the VemP-3 residues (146–156) in the ribosomal tunnel, hereafter denoted as *VemP-3R*. For each RNC, we prepared two initial conformations with either the A or B substate of U2506 (Fig. 1a).

Each RNC complex was placed within a rhombic dodecahedron periodic box of water ensuring a minimal distance of 1.5 nm between the solute and the box surface. Structural K^+^, Mg^2+^, Zn^2+^, and Cl^−^ ions were taken from a previous cryo-EM structure (PDB 5AFI)^27^ by superimposing the large and small ribosomal subunits separately. Excess ions were added at random positions at their respective concentrations in the cryo-EM experiment (100 mM KCl, 10 mM MgCl_2_) and, subsequently, the whole box was neutralized by adding sufficient K^+^ ions. The resulting simulation box contained about 2.1 million atoms.

In addition, isolated VemP in solution was studied by MD. The initial VemP conformation was taken from the cryo-EM model so one helix (VemP-2, VemP-3) or two helices (VemP-1) were present. The box size was chosen such that the minimal distance between the VemP and the box surface was 2.7 nm. For instance in VemP-1 case, this box size allows sampling sampling conformations with end-to-end distance lower than 8 nm. Note that the end-to-end distance of the fully extended VemP-1 conformation is 13 nm. However, our test simulations initiated from the fully extended conformation revealed that VemP collapses quickly (*<* 50 ns, Fig. S1) to a conformation with the diameter of about 3 nm, so the use of the smaller box size to sample collapsed conformations is justified.

For the simulations of the RNC, we used the Amber ff12SB force field^28^ for the solute with the parameter set of non-canonical nucleotides based on ff99SB,^29^ SPC/E water model,^30^ and Joung and Cheatham ion parameters. ^31^ This setup has proven reliable in several simulation studies including those of the ribosome. ^15,32^ For VemP constructs in solution, we used two force-fields of different families. First, the identical force field as for the RNC complexes, namely ff12SB and SPC/E. Second, we used a recent force field CHARMM36m,^33^ which was designed to better capture the equilibrium between folded and unfolded states. This protein force field was combined with the CHARMM-compatible TIP3P water model. ^34^ Table 1 summarizes the simulated systems.

**Table 1:**
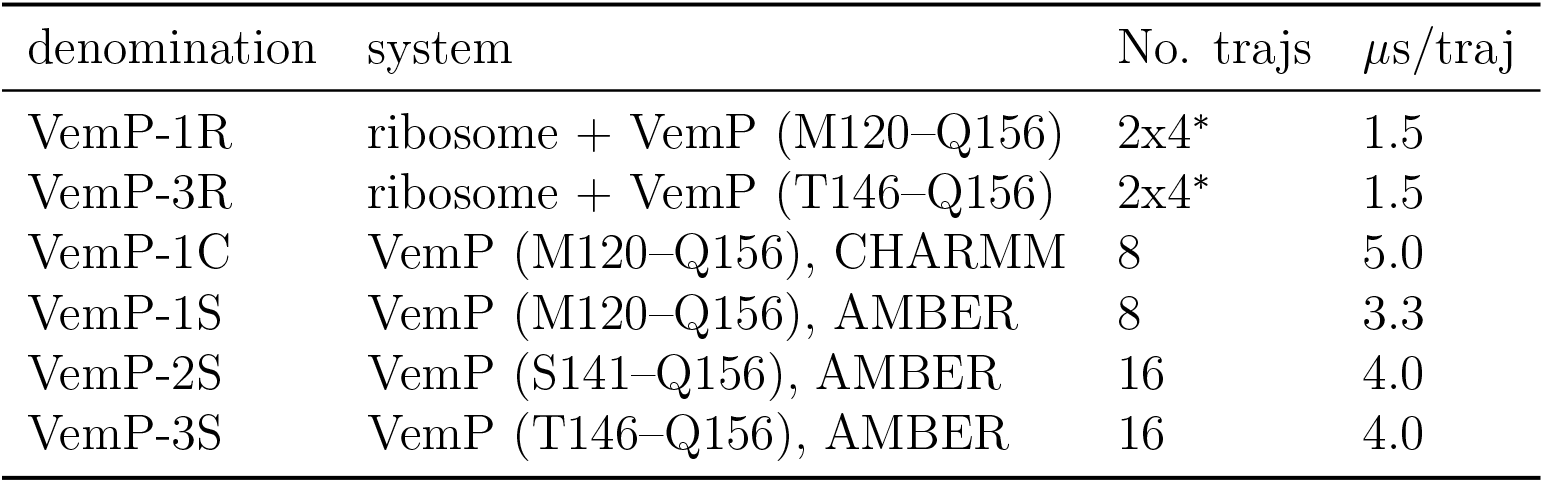
Summary of MD simulations. ^∗^Four trajectories for each of the A or B substate of the U2506 nucleotide.

| denomination | system | No. trajs | $\mu\text{s}/\text{traj}$ |
| --- | --- | --- | --- |
| VemP-1R | ribosome + VemP (M120–Q156) | 2x4* | 1.5 |
| VemP-3R | ribosome + VemP (T146–Q156) | 2x4* | 1.5 |
| VemP-1C | VemP (M120–Q156), CHARMM | 8 | 5.0 |
| VemP-1S | VemP (M120–Q156), AMBER | 8 | 3.3 |
| VemP-2S | VemP (S141–Q156), AMBER | 16 | 4.0 |
| VemP-3S | VemP (T146–Q156), AMBER | 16 | 4.0 |

### Molecular Dynamics Simulations

Each simulation system of Tab. 1 was equilibrated in several steps. The solvent was energy-minimized in roughly 25,000 steps keeping the solute fixed using Cartesian position restrains with a force constant of 1000 kJ mol^−1^ nm^−2^. Then during 500 ps of MD simulation, the solvent was heated from 10 K to 310 K using the v-rescale thermostat,^35^ while the solute was kept at 10 K. The inital velocities were selected randomly from a 10-K Maxwell-Boltzmann distribution. In the following 1 ns, the pressure (and the density of the simulated system) was equilibrated using the Berendsen barostat^36^ targeted to 1 bar. Then the solute was heated to 310 K and the position restraints were gradually released during another 20 ns MD simulation. Finally, the production runs were carried out at 310 K and 1 bar (*in vivo* conditions^10,16^) using the v-rescale thermostat and the Parrinello-Rahman barostat,^37^ respectively.

In all simulations, the long-range electrostatic interactions were computed by the particle-mesh Ewald algorithm^38^ with a 1.0-nm direct-space cut-off, interpolation order of 4 and 0.12 nm grid spacing. Van-der-Waals interactions were described by the Lennard-Jones potential with 1.0 nm cut-off. All bond lengths were constrained by LINCS ^39^ and the hydrogens were described by virtual sites^40^ to remove the fastest degrees of freedom, thus allowing for a time step of 4 fs (production runs).

All simulations were carried out using the GROMACS package^41^ version 2016 (RNC complexes) and 2020 (VemP constructs in solution).

### Circular Dichroism

The VemP constructs (VemP-1, VemP-2, and VemP-3) were synthesized by Biomatik using solid state synthesis at a 95% purity level. The lyophilized peptides were controlled by quantitative amino acid analysis and high-performance liquid chromatography. The peptides were then dissolved in buffer solutions containing 50 mM sodium-fluoride (NaF) and 10 mM sodium-phosphate (NaPi) buffer (pH = 7.2) as well as 0, 20, or 40 volumetric percent of 2,2,2-triflouroethanol (TFE). Final peptide concentrations were determined by measuring UV absorbance spectra in the 200-350nm range, and using the extinction coefficient at 280 nm, and ranged between 15–90 *µ*M.

The CD spectra of VemP constructs were measured at 25 °C using a Jasco J-815 Circular Dichroism Spectropolarimeter. The CD spectra were recorded between 180–260 nm at every 0.5 nm for both the peptides and blank buffer solutions in 1 mm cuvettes. The peptide CD spectra were baseline corrected, and normalized for protein concentration, cuvette path length, and the amino acid numbers to compute mean residue ellipticities (deg cm^2^ dmol^−1^). All CD spectra are shown in 1000 mean residue residue ellipticity units (kMRE). The CD spectra were truncated to 190–260 nm wavelength range to ensure a linear concentration-dependent signal intensity at all remaining wavelengths. CD spectra were analyzed for secondary structure content as described in Analyses.

### Analyses

The structures of exit-tunnel walls and VemP seen in the MD simulations were characterized by residue-wise root-mean-square deviation (RMSD) calculated between the MD conformations averaged over each of the trajectories and the conformations found in the cryo-EM model. Also for each VemP residue, we calculated the root-mean-square fluctuation (RMSF), which reports on how flexible/mobile each residue is within the MD ensemble. The details are provided in the Supplementary Information.

MD-derived structural RNC models were obtained at three levels of averaging. First, as a mean structure over each of the individual trajectories, second, by averaging over all trajectories initiated from the same U2506 substate (A or B), and third, by averaging over all RNC trajectories.

The SS of VemP was determined by the DSSP algorithm.^42^ This algorithm defines 8 secondary structures based on the overall geometry and hydrogen bonding patterns. All trajectories were processed by the GROMACS tool do dssp.

The SS of VemP constructs were also determined from measured CD spectra using the Bayesian estimation module (SESCA bayes.py) from the SESCA analysis package version 0.95. ^43,44^ The SS estimates were obtained using the DSSP-TSC1 basis set, which contains three secondary structures and six side-chain correction basis spectra. Side-chain corrections were determined based on the amino acid composition of the corresponding VemP construct. The SS estimates were calculated from probability-weighted ensembles of 100 000 SS compositions generated by 100 Monte-Carlo chains started from randomly chosen SSs.

## Results and Discussion

### The VemP conformation found in the ribosome is unstable in solution

The reasons why VemP folds into the specific secondary structures of two helices (residues 120–130 and residues 146–156) and the S-shape loop (residues 131–145) in the ribosome exit tunnel are not entirely clear. Su *et al*. speculated^16^ that the PTC likely evolved “to generally disfavour excessive secondary structure formation”. The preference for *α*-helices may be encoded in the VemP amino-acid sequence, it may be driven by the tunnel walls due to specific interactions between the peptide and the tunnel, in a nonspecific manner (*e*.*g*., by confinement or hydrophobicity of the environment), or there may be a combination of several factors.

As a first test of the hypothesis that VemP sequence has intrinsic preference for *α*-helices, we used two common SS prediction algorithms (Fig. 2a). The neural-network based three-state predictor JPred4^45,46^ classifies residues 122–134, which overlap with the residues of the outer helix, as helical (H), but only with a low confidence. In contrast, residues 148–153, which constitute the inner helix in the ribosome, are predicted as extended strand (E) with a high confidence. The NPS@ web server ^47^ yields a four-state prediction as consensus of 10 algorithms. In VemP, an extended strand (E) is predicted for residues 131–133 and 148–153, whereas remaining residues are predicted unstructured. To this end, the SS predictions provide no convincing evidence about sequence preferences of VemP to form the SS observed inside the ribosome.

**Figure 2:**
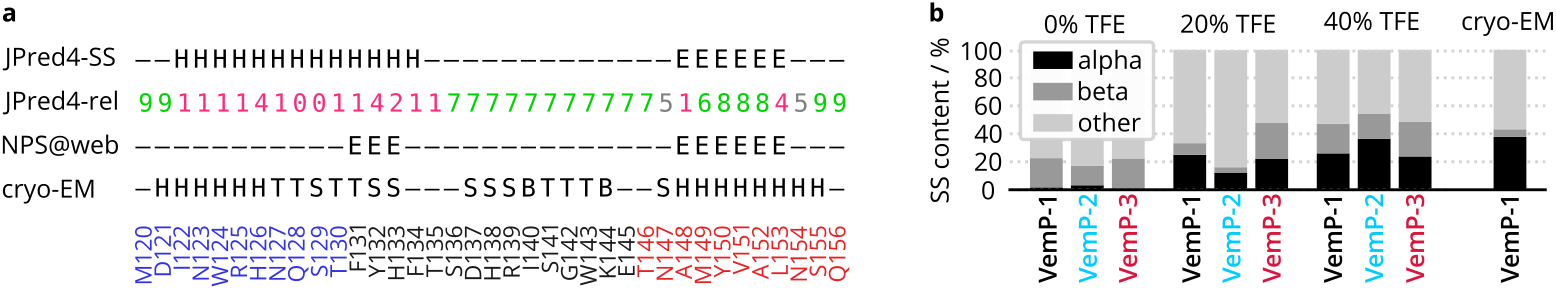
Secondary structure (SS) characteristics of VemP constructs. **a)** Sequence-based SS predictions of VemP-1 derived by JPred4^45,46^ (H=helix, E=strand) with the reliability values (JPred4-rel, larger is more reliable) and NPS@web^47^ (E=strand) and structure-based classification by DSSP ^42^ for the cryo-EM model (H=*α*-helix, T=turn, B=*β*-bridge, S=bend). The VemP sequence is color coded (blue=outer helix, red=inner helix). **b)** SS content of three VemP constructs as inferred by SESCA analysis of CD spectra in three solvents differing in 2,2,2-trifluorethanol (TFE) molar fraction. For comparison, the SS content of the VemP cryo-EM model ^16^ in the ribosome tunnel is shown as classified by DSSP.

To obtain quantitative SS propensities, we carried out CD measurements in solvents with different hydrophobicity (spectra in Fig. S6) and analyzed them by SESCA.^43,44^ In water, none of the VemP constructs shows a sizeable helical content (Fig. 2b). In fact, each VemP construct appears to be about 80% disordered, and partially adopts extended or *β*-strand-like conformations. Increasing the hydrophobicity of the solvent by adding TFE (20 mol.%) in the CD measurements resulted in an increased estimated helical content. For VemP-1 and VemP-3, the helical content is about 20%, for VemP-2 it is about 12%. After increasing the hydrophobicity of the solvent even further (40 mol.% TFE), the helical content increased in VemP-2 further to 36%, while the helicity of VemP-1 and VemP-3 remained similar to their helical content in the lower TFE concentration.

The helical content of VemP in the exit tunnel can be estimated from the cryo-EM model as a ratio of the number of helical AAs classified by DSSP and the total number of AAs (Fig. 2a). For VemP-1 in the ribosome tunnel, the helical content is 38%. The CD spectra indicate that VemP-1 in solution has a helical content of close to 0% for water and about 25% for the two hydrophobic solutions. This finding suggests that the hydrophobicity of the VemP environment alone does not suffice to explain the SS content in the ribosome.

To understand the helicity loss of the VemP conformation found in the ribosome tunnel, and the SS preferences of VemP components, we carried out atomistic simulations of VemP fully solvated in water. For all constructs, the simulations were initiated from their conformation in the ribosome. The DSSP classification reveals notable helical regions in all constructs (Fig S3). On average, the helical content of VemP-1 remains roughly constant in the Amber force field (VemP-1S in Fig. 3) or decreases from about 50% to 25% in the CHARMM force field (VemP-1C in Fig. 3). VemP-3 is the only construct studied that shows a substantial decrease of the helical content (from 50% to about 10%) within the simulation time span.

**Figure 3:**
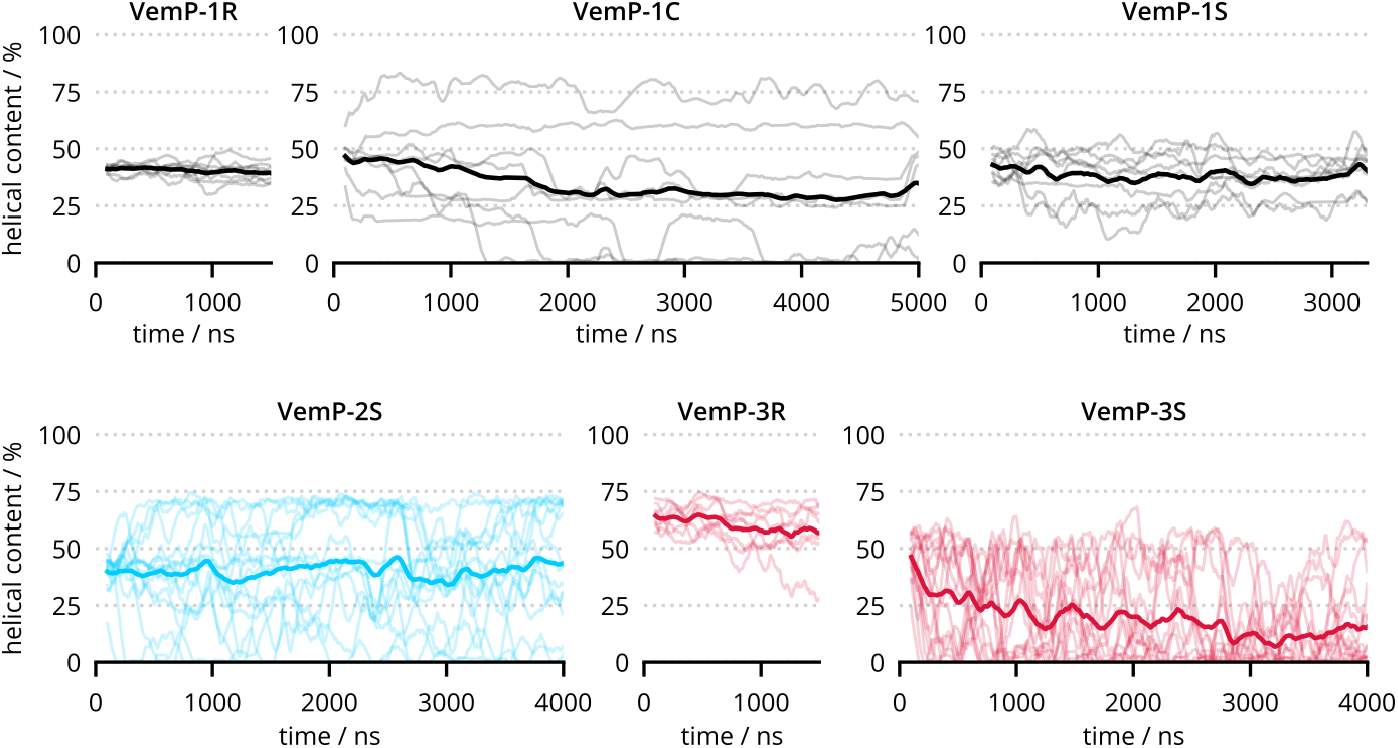
Helical content of VemP-1 (black), VemP-2 (cyan), and VemP-3 (red) calculated from MD simulations in the ribosome (R) and in solution (C and S, two force fields). In each panel, individual trajectories are shown as thinner lines, while their average is shown in the thicker line.

We further analyzed the helical content of each VemP residue (Fig. 4). In the ribosome, two regions of VemP are helical, residues 120–130 (outer helix) and 146–156 (inner helix). The helical content of these two regions decreases irrespective of the construct or force field. The most pronounced decrease is observed in VemP-3S, and for the inner helix in VemP-1C. However, residues outside the two regions, which are non-helical in the ribosome and were non-helical in the beginning of MD simulations in water, increase their helical content slightly. This results in an overall peptide helical content roughly constant or decreasing slowly (Fig. 3).

**Figure 4:**
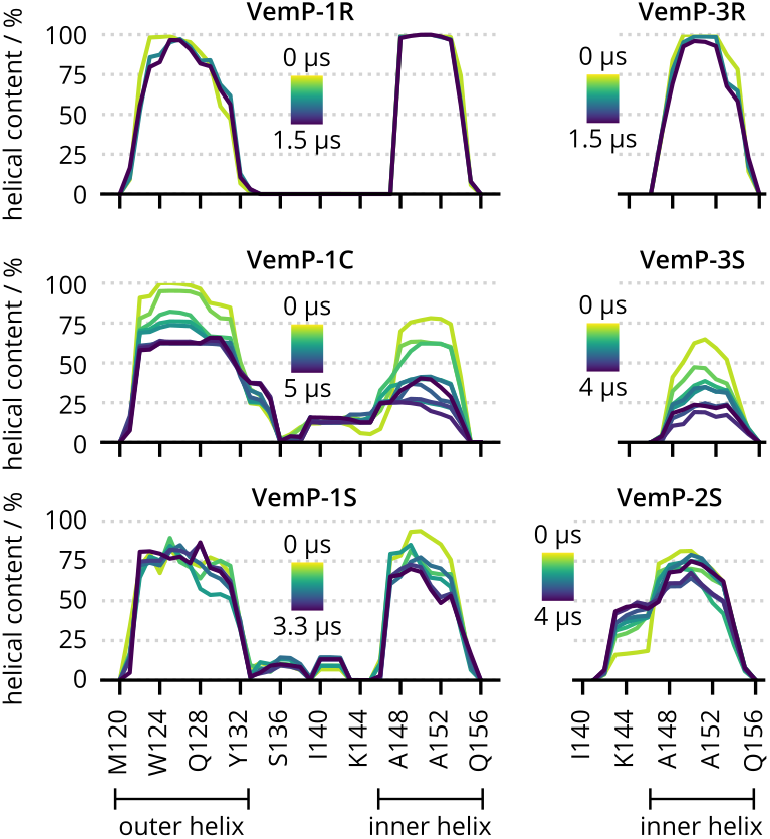
Residue-wise helical content of VemP-1, VemP-2, and VemP-3 calculated from MD simulations in the ribosome (R) and in solution (C and S, two force fields). The helical content was calculated for several time blocks colorcoded from yellow (beginning of simulations) to navy (end of simulations) and averaged over independent trajectories. The lines are separated in time by 500 ns; longer simulations resulted in more lines.

The helical content does not converge during the simulation time as can be seen by the differences in the individual trajectories (Fig. 3). As the CD measurements indicate, the helical content in water should drop to close to zero eventually. The fact that this did not happen during the several microseconds simulation time suggests that the *α*-helical VemP conformations represent a relatively deep local free-energy minimum. Here, imperfections of the used force fields are certainly a factor, as reflected in the different kinetics of helicity changes in the simulations.

Taken together, our results show that VemP requires an environment different from bulk water to adopt stable helical conformations. To evaluate the role of the exit tunnel in VemP helix formation, we compared the conformational ensembles obtained from MD simulations of VemP in solution with MD ensembles inside the ribosome.

In the ribosome, both helices remain fully helical on the simulation time scales (*µ*s) and close to their conformations in the cryo-EM structure, as indicated by the backbone RMSD below 0.14 nm (Fig. S4). The overall helical content is about 45% and remains constant throughout the simulations (Fig. 3). Much smaller variations of helical content among individual trajectories, when compared to the simulations in water, concur with the stabilizing effect of the ribosome tunnel.

The residue-wise helical content obtained from the MD simulations in the ribosome agrees well with the cryo-EM structure classification (VemP-1R in Fig. 4). In the simulations, residues 120–122 seem to be less helical compared to the cryo-EM model. This is probably due to the fact that residues 26–119 were not included in the simulations, whereas they were part of the VemP construct used for cryo-EM experiments. ^16^ The residues up to 119 were not resolved in the cryo-EM model, likely because they are more flexible than the C-terminal residues, but their presence possibly stabilized the VemP N-terminal helix.

The tunnel walls near the PTC are formed partly by rather hydrophobic rRNA nucleobases. In addition, the negatively charged sugar-phosphate backbone of the rRNA also contributes to the peptide-tunnel interactions. The constriction site is also highly hydrophilic due to the positive side chains of uL4 and uL22 which point into the tunnel. Petrone *et al*. reported^48^ that the exit tunnel disfavors hydrophobic amino acids which would suggest poor interactions between hydrophobic NCs and tunnel walls. Moreover, the exit tunnel is filled with water whose behavior differs from bulk, in particular in dielectric and diffusion characteristics. Lucent *et al*. argued^49^ that the water behavior favors compaction of hydrophobic sequences. This goes in line with the CD measurements in TFE solution, where VemP shows a higher helical content than in water. Still, the helical content of VemP-1 is much lower in solution (*<*26%) than in the ribosome tunnel (about 45%). Thus the hydrophobicity cannot fully explain the SS structure observed in cryo-EM. Especially the stability of the inner VemP helix seems to require additional explanation, other than the hydrohpobicity effect alone. The CD spectroscopy in TFE indicates the helical content at most 24% while in the ribosome tunnel this portion of VemP has helical content of 80%. Hence in the next part, we focus on the specific interactions between VemP and tunnel walls identified previously by cryo-EM.^16^

### Analysis of U2506 substates reveals a structurally conserved VemP region

The cryo-EM model represents the average of A and B substates with the exception of U2506, where the electron densities are different enough to be resolved as two states. In contrast, MD simulations allow sampling RNC conformations separately, depending on the initial substate of U2506. Indeed, we observe two distinct conformational ensembles of VemP and its neighborhood for the A and B substates. During the 1.5 *µ*s of simulations, no transition between the U2506 substates happened. Differences between the A and B ensembles are found up to 2 nm away from the C-terminus of VemP, or to a distance of roughly three nucleotides from the U2506.

Notably, in the two substates, the position of the VemP C-terminal residues Y150–Q156 differs (red arrows in Fig. 5de). In substate A, U2506 partially occupies the ribosomal A site, and it pushes the helix towards the tunnel exit more than in substate B. There is also an interaction between N154 and U2506 in substate B (but not in A), which can influence the positioning of the VemP C-terminus. The average positions of the C-terminal Q156 in the direction of the exit tunnel differ by about 0.15 nm between the two substates.

**Figure 5:**
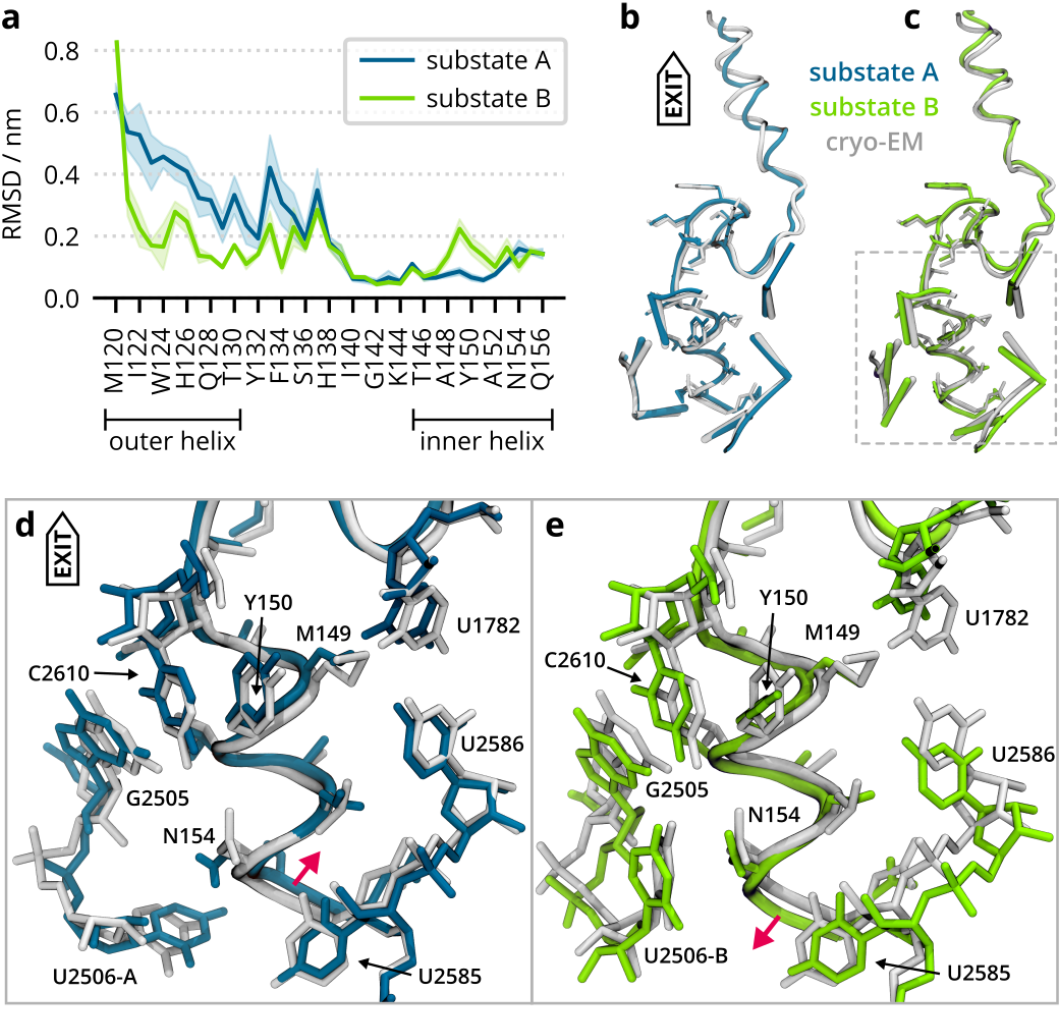
Structural differences between A and B substates. **a)** Residue-wise RMSD between the cryo-EM model and structures averaged over each of the trajectories. The mean RMSD value over the 4 independent trajectories is shown as a line, the shaded area represents the standard error of the mean. **b)** Comparison of the atomistic model obtained as an average over all trajectories initiated from U2506 in substate A (in teal) and the cryo-EM model (in gray). Selected nucleotides are represented as sticks, whereas the side chains of residues 143–156 are shown as licorice. **c)** Same as b) but for the substate B. **d)** The detail of the VemP inner helix and its surroundings as framed in c), but depicted as licorice. The MD-derived model from trajectories initiated from the substate A is is in teal, cryo-EM in gray. Red arrow indicates a dislocation of the helix with respect to the cryo-EM model. **e)** Same as d) but for the substate B.

To quantify the structural differences between A and B substates, we calculated residue-wise RMSDs of non-hydrogen atoms between the average conformations found in MD and the cryo-EM model and analyzed MD-derived atomistic models (Fig. 5).

The plot of residue-wise RMSDs (Fig. 5a) has two main features. First, the N-terminal part up to H138 has a higher RMSD in substate A than in B, showing that the structures of the outer helix found in the four independent trajectories deviate from the cryo-EM model more for substate A than for substate B. However, averaging the structure over all trajectories of the same substate, the agreement with cryo-EM is good in both substates (Fig. 5bc). Thus the discrepancies can be attributed to the missing N-terminal part up to M120, which probably further stabilizes the outer helix within the tunnel. The N-terminal part was present in the cryo-EM experiments, but is missing in the simulations. Second, a portion of the inner helix (A148–A152) has higher RMSDs in substate B than in A. The structural variations are observed for AA side chains rather than the VemP backbone. The largest difference is found for M149 with a distance of 1.1 nm from the VemP C-terminus. The U2506 substate affects the vicinity of M149, namely the hydrogen bonding between U1782 and U2586, and stacking of Y150 with C2610. The former interaction is affected through U2585, while the latter through G2505 (Fig. 5de).

Importantly, our observations are consistent with the cryo-EM model. When we average all RNC trajectories (A and B), the structural agreement between MD and the cryo-EM model is excellent (Fig. 6a). The backbone RMSD of VemP C-terminal R139– Q156 is only 0.05 nm. Both backbone RMSD of the entire VemP as well as the non-hydrogen atom RMSD of selected nucleotides around the inner helix (apart from U2506) are below 0.1 nm.

**Figure 6:**
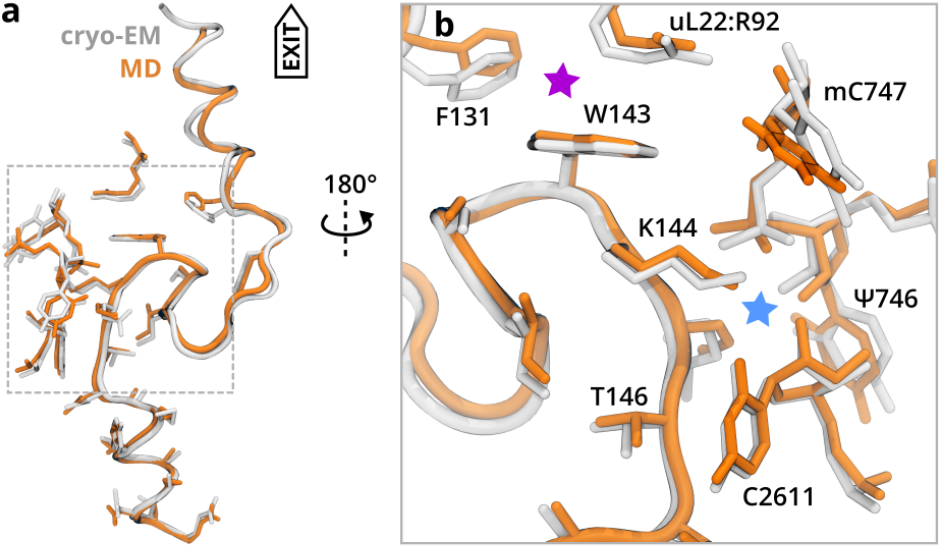
**a)** The MD-generated model (in orange) was obtained as an ensemble average of all RNC trajectories (A and B). The cryo-EM model is in gray. **b)** Context of the anchoring VemP part. W143 stacks to R92 of ribosomal protein uL22 and is involved in a hydrophobic contact with F131 of VemP (magenta star). K144 side chain directs to a pocket formed by the phosphate between pseudouridine 746 and 5-methylcytosine 747 and with oxygen of C2611 (blue star).

Intriguingly, there is a region between I140 and K144, or even N147, which is structurally very well conserved, independent of the U2506 substate the MD simulation was started from. These residues are located on the N-terminus of the inner helix near the constriction site. The group of residues includes W143 and K144, which were identified as critical for stalling. ^10,21^ They feature among the lowest RMSDs due to several interactions with the tunnel wall (Fig. 6b). The interaction of the positively charged side chain of K144 with the negatively charged phosphate of mC747 appears to be crucial for stalling, as reported by previous biochemical experiments. The stalling was alleviated by any mutation of K144 apart from positively charged AAs (R, H). ^21^ Likewise, W143 forms a hydrophobic contact with R92 of uL22 and F131 of VemP. This is also the place, where the outer helix may play a role in translational arrest as suggested by Su et al., ^16^ additionally stabilizing the VemP region around W143.

It is conceivable that due to the incremental elongation of NCs, these residues act as an *anchor*, which slows down the VemP progress through the tunnel. This may modulate the free energy landscape of the subsequent VemP intermediates such that the deepest minimum corresponds to the *α*-helix, which inhibits the peptide bond formation eventually. A mechanism of this kind was recently reported by Valle et al. in the context of ornithine sensing by ribosomes. ^7^ In their system, a NC sensing domain binds the tunnel around U464 located about 4 nm from the PTC (not shown), whereas in the case of VemP the anchoring ribosomal residues are around Ψ746 and C2611 (Fig. 6b), which is about 3 nm from the PTC.

We also inspected the flexibility of VemP and its surroundings through the residuewise RMSFs and their projections onto the cryo-EM structure (Fig. 7). Similar to the pattern observed for RMSDs, the outer helix is more flexible that the inner helix. This is consistent with the exit tunnel diameter, which is smaller near the inner helix and PTC, and larger near the outer helix beyond the constriction site. MD simulations also show that the flexibility of the outer helix is not much sensitive to U2506 substate. In contrast, the region of VemP near M149 shows higher RMSFs for substate B than for A (alike to RMSDs, Fig. 5). RMSFs of the VemP residues near K143 are among the lowest, which brings another piece of evidence about the large structural conservation of the VemP anchor.

**Figure 7:**
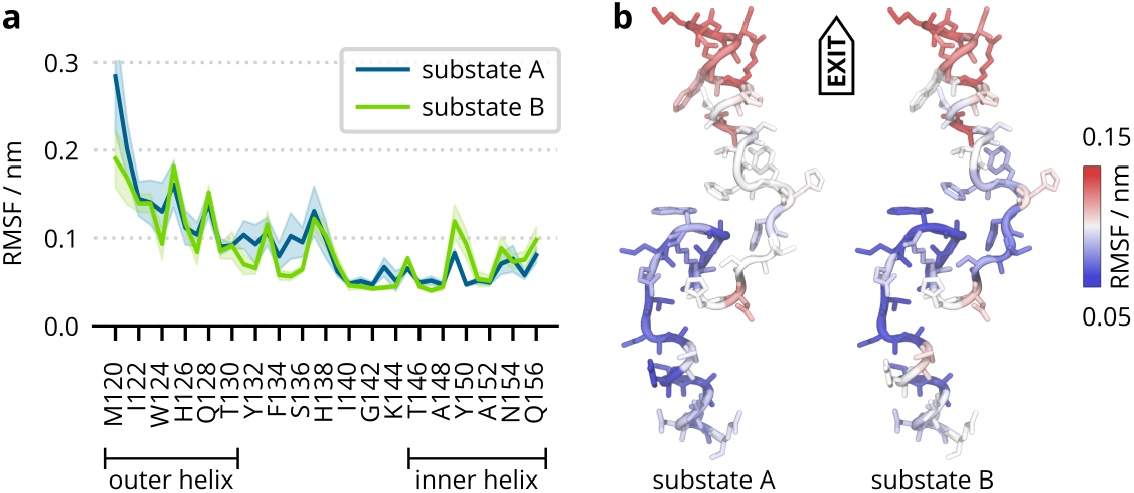
**a)** The residue-wise root-mean-square fluctuation (RMSF) calculated from trajectories started with U2506 in substates A (black) or B (green). The mean value over 4 independent trajectories is shown as line, the shaded area represents the standard error of the mean. **b)** Projection of the RMSF from MD onto the cryo-EM model. The color scale goes from blue (rigid), through white to red (flexible).

### The inner VemP helix is stabilized in the ribosomal tunnel

To assess the role that the N-terminal VemP parts (S-shape loop and outer helix) play in the stalling mechanism, we simulated VemP-3 (inner helix) in the ribosome tunnel. In solution, the helical content of VemP-3 is very low in both CD spectroscopy and simulations, so it is conceivable that the helix may also unfold in the tunnel without being held in place by the anchor and loop. Because the helix sterically prevents U2506 and U2585 from adopting translation competent conformations, unfolding of the helix could release the stalling.

We observe that throughout all 8 VemP-3R simulations the inner helix remains helical regardless of the initial position of U2506 as reported by the per-residue helical content (Fig. 3) and backbone RMSD (Fig. S5). However, the helical content is lower than in VemP-1R, in particular for the two N-terminal AAs (T146, N147). Recall that the starting conformation of VemP-3R was always helical as found in the cryo-EM. One of the 8 independent simulations shows a decrease of helical content from about 60% to 25% within the simulation time. This hints that the stability of the inner helix alone in the tunnel (VemP-3) is lower than the stability of the inner helix when the outer helix is present (VemP-1).

There is a notable difference in the VemP-3 helix position relative to the tunnel between the A and B substates. While the inner helix stays is place in substate A, in substate B the entire helix shifts by about 0.15 nm towards the tunnel exit (Fig. 8). The observed VemP-3 shift resembles a spring release triggered by the *in-silico* removal of the large N-terminal part. This suggests that the stalled VemP peptide is under strain in the narrow space near PTC and that the strain is more pronounced when U2506 is in state B, where it directs towards the tunnel exit and further reduces the diameter of the tunnel near PTC.

**Figure 8:**
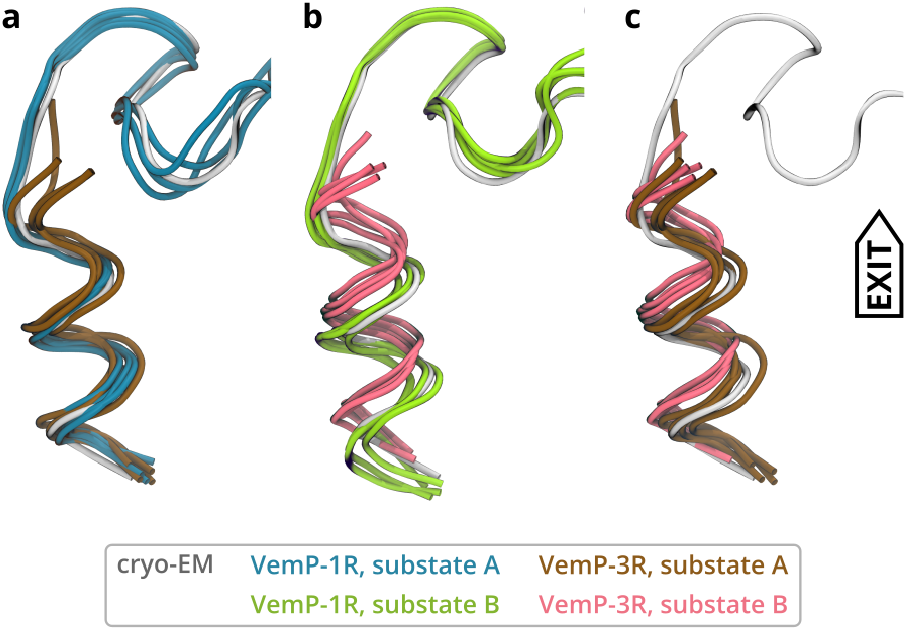
Comparison of VemP-1 and VemP-3 in substates A **a)**, and B **b)** in the ribosomal tunnel. VemP-3 substates are compared in **c)**. The cryo-EM model is shown in gray for reference. Each structure represents the mean conformation of one of the four independent trajectories.

The nucleotides surrounding the helix C-terminus adopt similar conformations as in the VemP-1R case. U2506 and U2585 stay unaffected by the missing S-shape loop and C-terminal helix in VemP-3R simulations. As expected, the nucleotides closer to the constriction site, which originally resided near the removed VemP part, changed their structure and dynamics. For instance, U1782 and C2609 were more flexible compared to the case of VemP-1R.

We expect VemP-3 to be stalling incompetent, although no biochemical characterization of VemP-3 has been done so far. It was reported that a slightly longer VemP construct with residues 138–156 shows a stalling efficiency reduced to 10% compared to the full-length VemP.^10,16^ Even though we do not see complete unfolding of the inner helix on the time scale of the simulations, the reduced helical content and the shift of the helix might be a first step towards the release of the U2506 and 2585. The faster and more complete unfolding observed in the simulations of VemP-3 in solution suggests that the interactions between the helix and tunnel constitute a free-energy barrier for helix unfolding.

## Conclusions

Nascent chains can stall ribosomal translation by various mechanisms which include the inhibition of tRNA accommodation, peptide-bond formation, tRNA translocation, and peptide release.^1^ A wealth of structural information on the conformations of stalling peptides bound to the ribosome lead to the identification of the molecular mechanisms underlying the inhibition of these steps of translation. However, general principles of how nascent chains adopt stalling conformations remain unclear. Here we studied the molecular details of translational arrest of the VemP peptide, which folds into two compact helices in the ribosomal exit tunnel and interferes with tRNA accommodation.

The sequence of VemP and even of its inner helix is unique and as for now it has been found only in the genus *Vibrio*. Thus it seems to be evolutionary highly optimized for its purpose.^21^ Still, the sequence itself lacks strong helical propensity in water, as we have shown by CD spectroscopy.

We found that the conformations and dynamics of VemP in solution are very different compared to VemP within the ribosomal tunnel; obviously it is to a large extent determined by the environment. We found that both the hydrophobicity of the environment as well as specific interactions between conserved residues of VemP and the environment contribute to the stability of the *α*-helix.

Additionally, a detailed analyses of our MD simulations of two U2506 substates revealed a region of VemP, which is structurally conserved and rigid. The results suggest two distinct roles, which the VemP AAs play during translation arrest. The inner helix (residues 146–156) clearly interferes with the two critical nucleotides,^16^ but it needs a portion of about 10 VemP AAs (I135–E145) with several specific and possibly strong interactions around W143 and K144 to become stable and well-localized within the tunnel. We argue that these residues serve as an anchor, which suggests a simple model that explains the sequence dependent (Fig. 9) formation of the inner VemP helix and the subsequent translational arrest.

**Figure 9:**
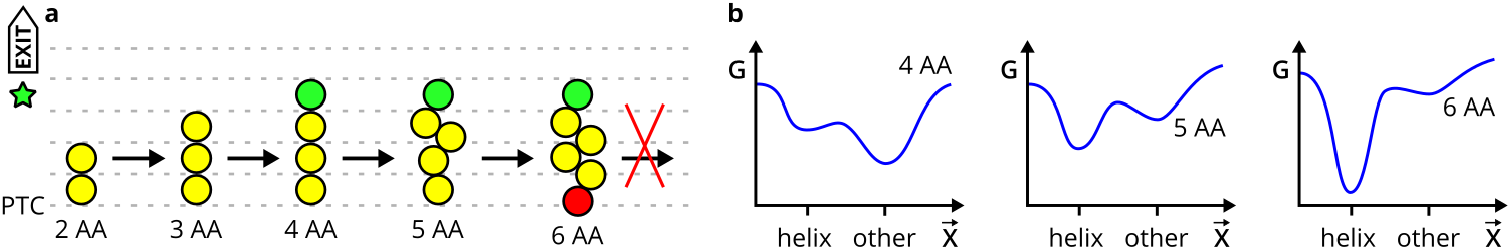
A model for anochor-mediated VemP translational arrest. **a)** Nascent peptide (NP) elongation proceeds until the anchoring amino acid (AA, green circle) reaches the anchoring point of the tunnel (green star). The anchoring interactions prevent the NP from translocating further, such that secondary structure content increases upstream until a sterical clash with the PTC (red circle) occurs, ultimately causing arrest. For simplicity, less AAs (yellow circles) than present in VemP is shown. **b)** Sketch of a possible conformational free energy (G) landscapes of VemP intermediates in the tunnel with respect to a generalized conformation coordinate *X-+* that characterizes NP translocation. The landscapes corresponds to NP from a) of 4, 5, and 6 AAs.

In this model, we assume that the rate of AA addition (in bacteria several AA per second) is much slower than the relaxation times and folding dynamics of the VemP peptide within the tunnel. Therefore during each prolongation step (snapshots in Fig. 9), the VemP peptide reaches local conformational equilibrium. VemP intermediates translocate through the tunnel until the anchoring AAs interact with the tunnel walls (Fig. 9a). The interaction changes the conformational free-energy landscape of the subsequent VemP intermediates (Fig. 9b). Our CD measurements suggest that the free-energy minimum corresponding to the *α*-helix is initially shallow and determined by non-zero sequence helical propensity in a hydrophobic environment. Upon every AA addition the minimum becomes deeper due to specific interactions of N-terminal VemP residues and tunnel walls, as determined by cryo-EM^16^ and observed in our MD simulations.

It remains unclear whether the outer helix is formed before of after the anchoring interactions are formed. Nevertheless, at least through the F131-W143 contact there is a mutual stabilization of the outer helix and the anchor, which could also explain the previously observed dependence of stalling efficiency on the presence of the outer helix.

The anchor-mediated translational arrest represents an allosteric mechanism that a local change of the interaction between amino acids and the anchor site causes a distant effect at the PTC. In fact, similar allosteric effects onto the PTC have been seen for other peptides such as ErmCL^15^ or SecM.^50^ The allostery here, however, differs from the classic view, where a conformational change within a *single* biomolecule or biomolecular complex is responsible for the signal transduction. ^51^ In our model, the signal is transferred through a modification of free-energy landscapes of multiple chemical reactions involving VemP intermediates.

The assumption that each VemP intermediate reaches conformational equilibrium can be expressed in terms of reaction kinetics that the rate of peptide bond formation is much lower than the rate of NP translocation through the tunnel. This suggests that the rate of NC translocation may be essential for the stalling mechanism, which would be in line with accumulated evidence that the rate of elongation has profound regulatory consequences. ^52^

## Acknowledgement

We thank Andrea C. Vaiana and Ting Su for valuable discussions and Roland Beckmann for pointing us to VemP.

## Authors’ Contribution

MHK, LVB and HG designed the research. MHK performed MD simulations. MHK, LVB, GN analyzed MD simulations. GN, JK, SMV performed and analyzed CD measurements. MHK, LVB, GN interpreted the results and wrote the manuscript with the input from other co-authors. All authors have read and agreed to the published version of the manuscript.

## Supplementary Information Analyses

### Residue-Wise Root-Mean-Square Deviation

The trajectories were superimposed onto the cryo-EM model using the backbone atoms of the exit tunnel (Tab. S2). The average structures were calculated for 1000–1500 ns of each of the trajectories. The RMSD_*s,t*_ of a residue *s* between conformation in the trajectory *t* and the cryo-EM model was calculated as

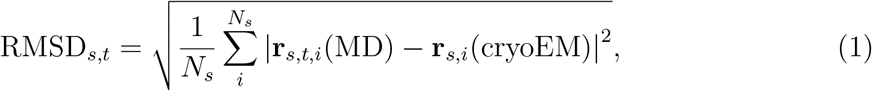

where **r**_**s**,**t**,**i**_ is the mean value of the position vector of the atom *i* in residue *s* over the portion of trajectory *t*, **r**_*s,i*_ if the position vector of the atom *i* in residue *s* in the cryo-EM model, || stand for the norm of a vector, and *N*_*s*_ is the number of atoms of residue *s*.

The RMSD of a residue *s* was then calculated as the mean value of RMSD_*s,t*_ of the trajectories.

### Root-Mean-Square Fluctuations

The root-mean-square fluctuation of the residue *s* over the trajectory *t* was calculated as

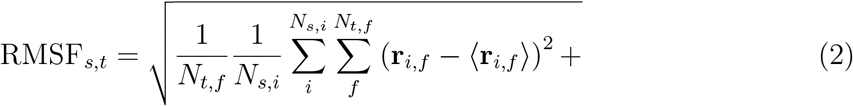

where **r**_*i,f*_ is the position vector of atom *i* in the trajectory frame *f*, the brackets ⟨ ⟩ stand for the mean value, *N*_*t,f*_ is the number of frames of trajectory *t* and *N*_*s,i*_ is the number of atoms in the residue *s*.

The RMSF of a residue *s* was then calculated as the mean value of RMSF_*s,t*_ of the trajectories.

## Tables

**Table S2:**
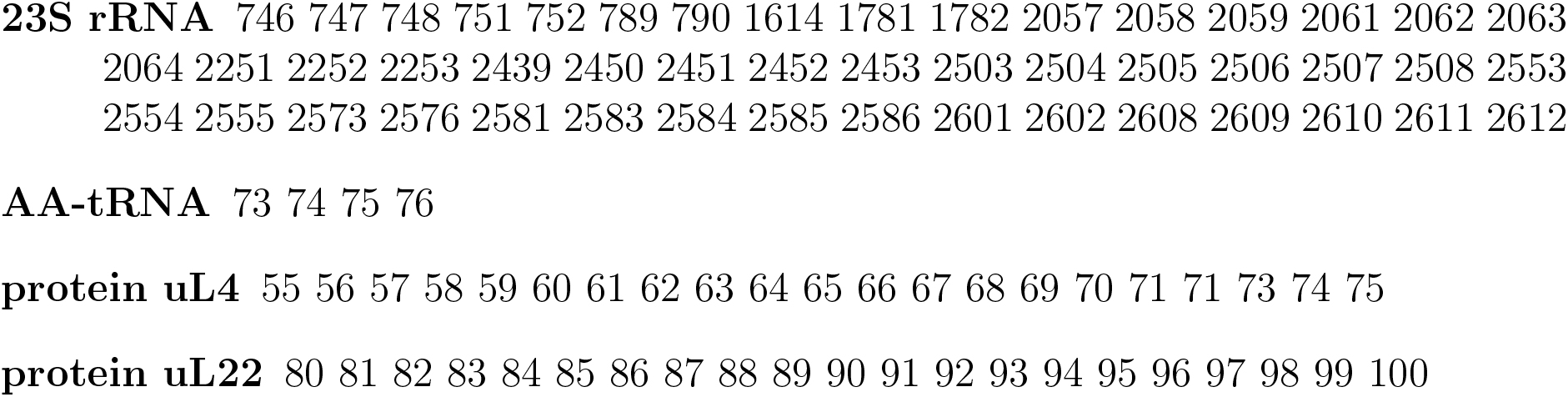
Numbers of the residues used for the least-square alignment of MD trajectories (*E. coli* numbering).

## Figures

**Figure S1:**
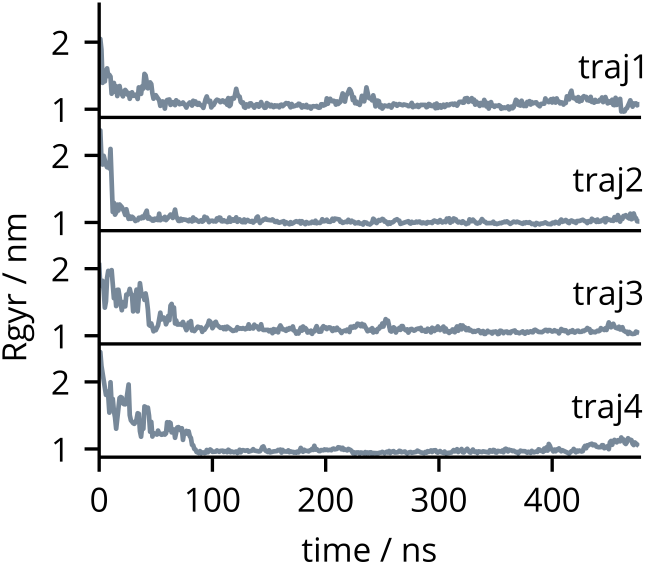
Radius of gyration of the VemP1 in solution (i.e. out of the ribosome) as a function of simulation time. The simulations were started from the fully extended VemP conformation.

**Figure S2:**
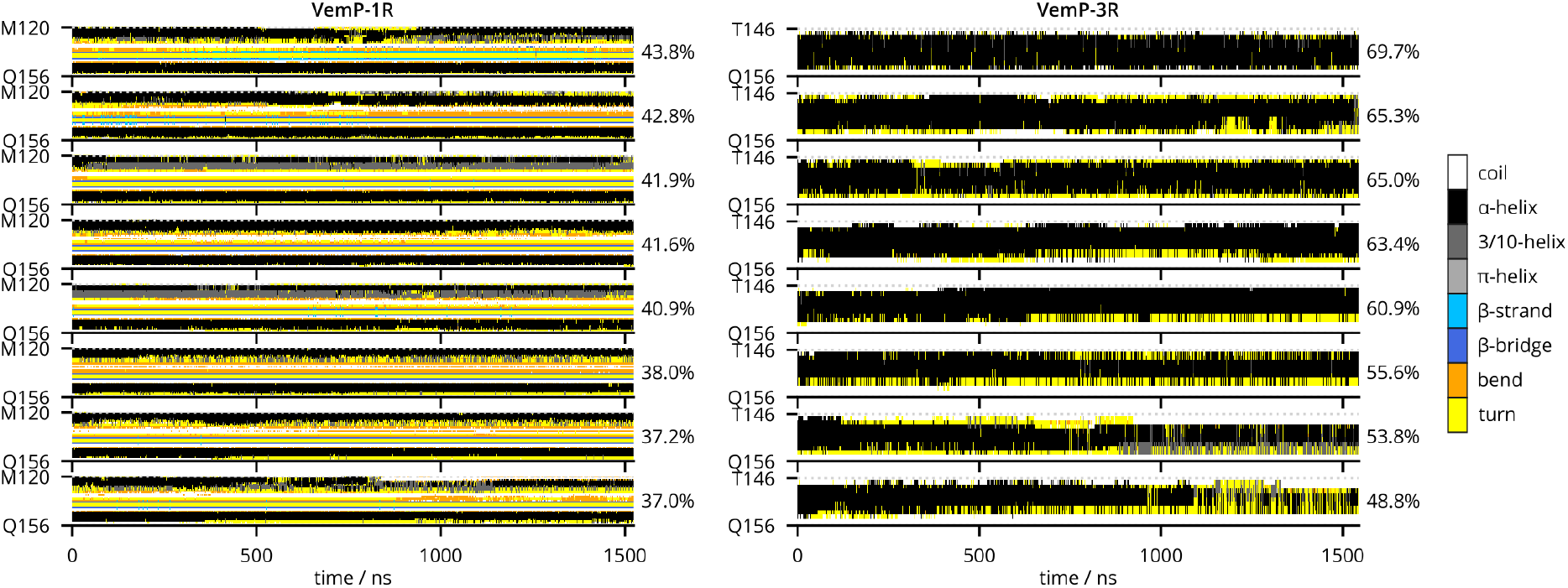
Secondary structure classification of VemP1-1 and VemP-3 in the ribosome tunnel as defined by DSSP algorithm. The timelines are sorted according to the overall helical content, which covers the *α*-helical and 3/10-helical motives and is shown on the right in %.

**Figure S3:**
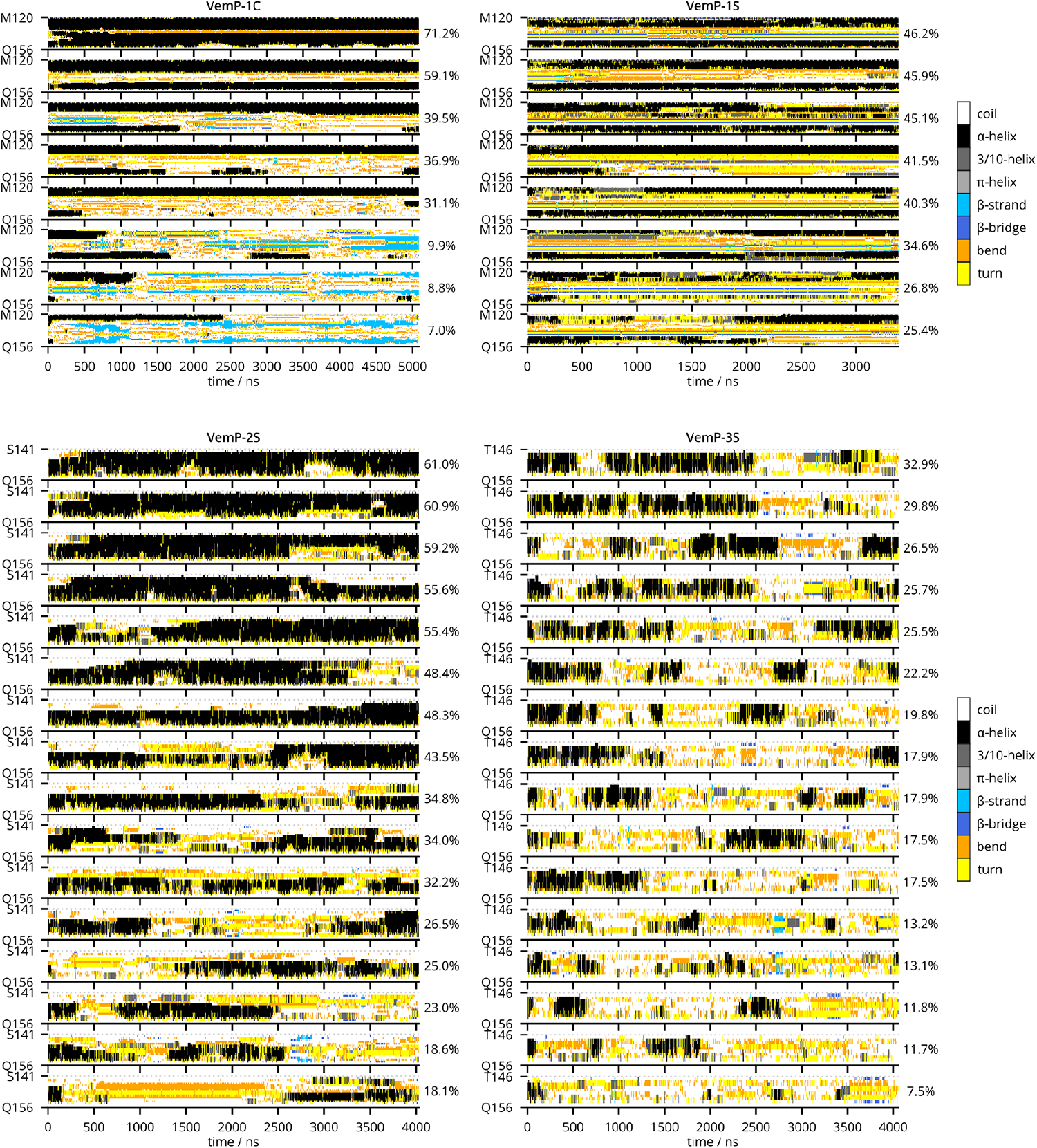
Secondary structure classification of VemP1-1, VemP-2 and VemP-3 in solution as defined by DSSP algorithm. The timelines are sorted according to the overall helical content, which covers the *α*-helical and 3/10-helical motives and is shown on the right in %.

**Figure S4:**
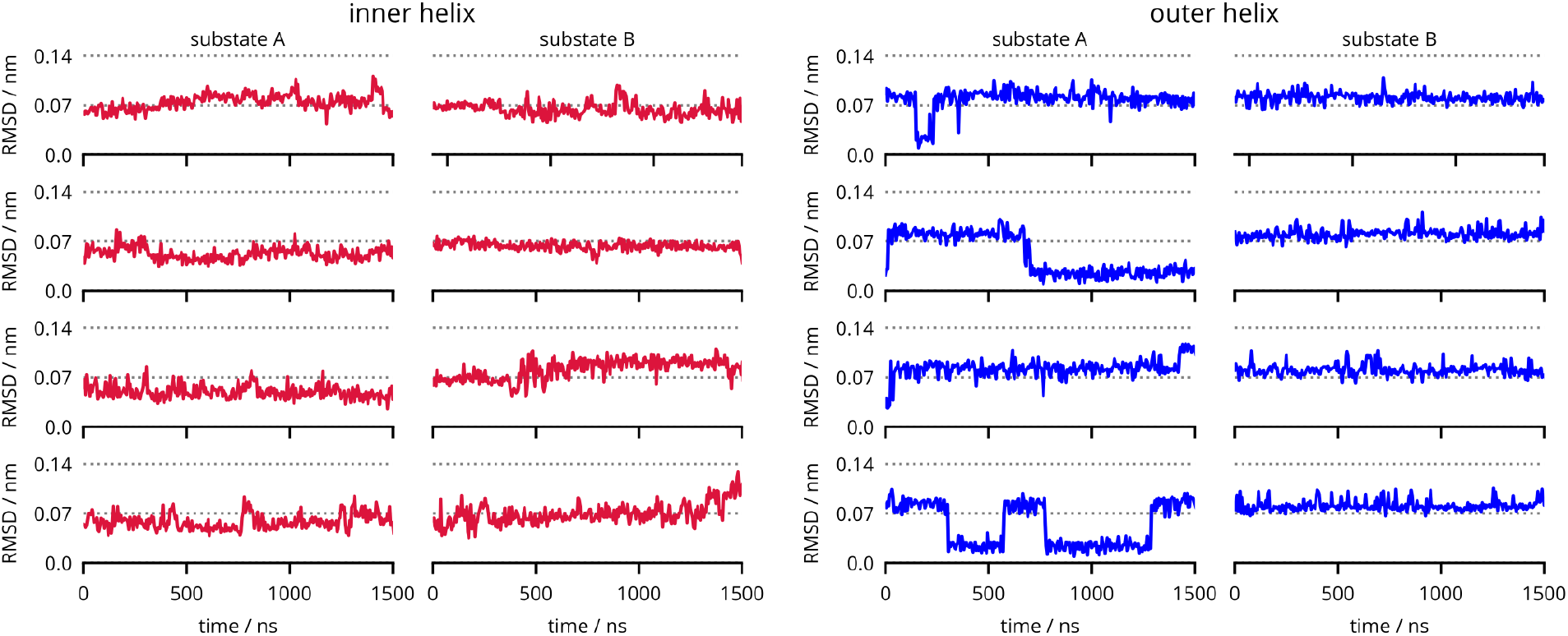
A backbone root-mean-square deviation (RMSD) of VemP helices in the ribosome tunnel after their least-square self-alignment. RMSDs were obtained in VemP-1R simulations.

**Figure S5:**
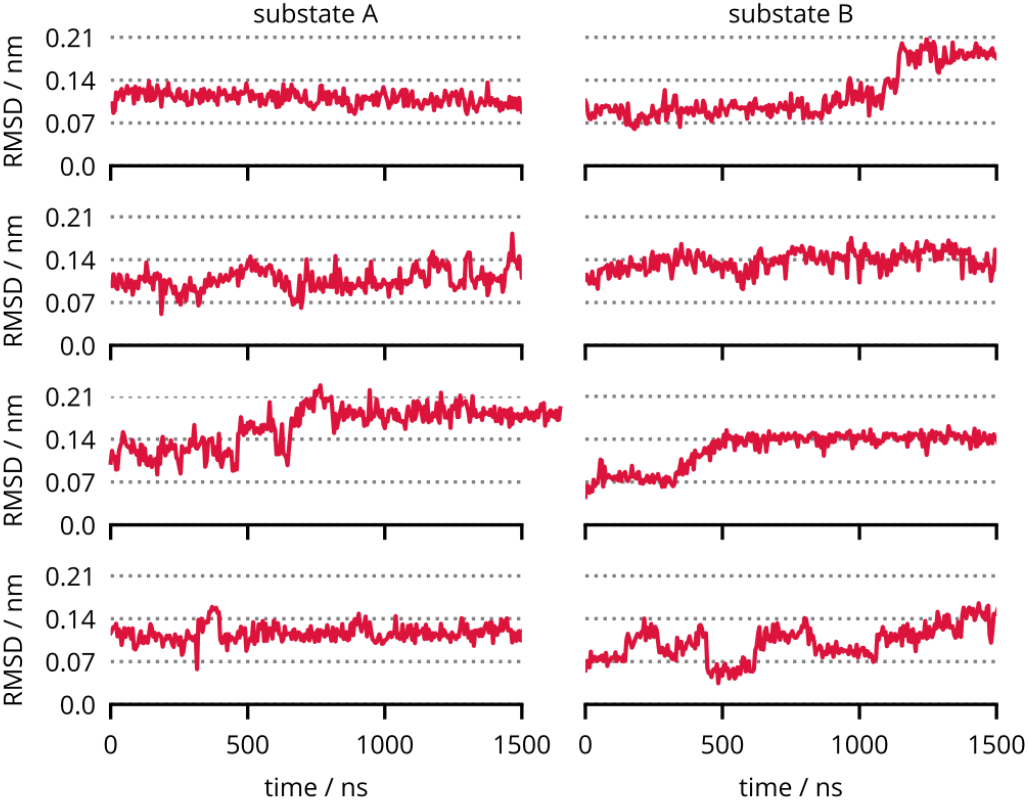
A backbone root-mean-square deviation (RMSD) of VemP-3 (inner helix) in the ribosome tunnel after the least-square self-alignment.

**Figure S6:**
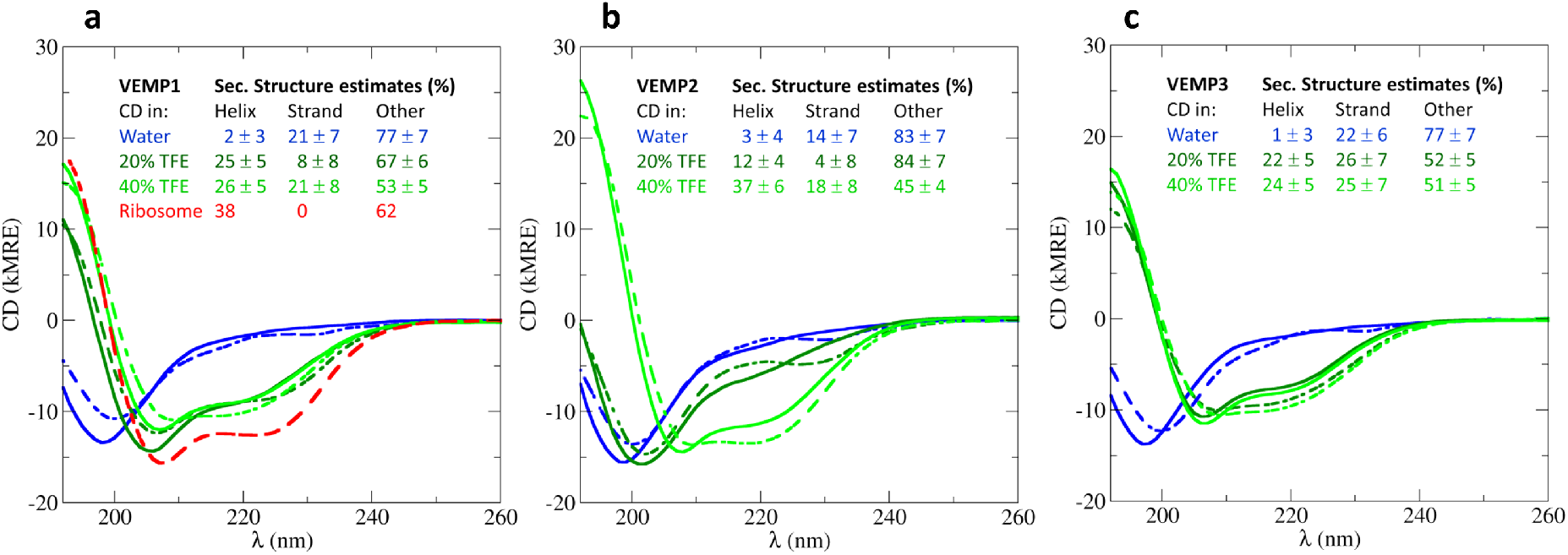
Measured CD spectra (solid lines) and CD spectra predicted from estimated SSs compositions (dashed lines) for VemP-1 (a), VemP-2 (b) and VemP-3 (c). The CD spectra predicted for VemP-1 in the ribosome (red dashed line) is calculated from the SS of VemP in the cryo-EM structure.

## References

(1) Ito; K., Chiba, S. Arrest peptides: cis-acting modulators of translation. Annual review of biochemistry 2013, 82, 171–202.

(2) Wilson, D. N., Arenz, S., Beckmann, R. Translation Regulation via Nascent Polypeptide-Mediated Ribosome Stalling. Current Opinion in Structural Biology 2016, 37, 123–133.

(3) Chandrasekaran, V.; Juszkiewicz, S.; Choi, J.; Puglisi, J. D.; Brown, A.; Shao, S.; Ramakrishnan, V.; Hegde, R. S. Mechanism of ribosome stalling during translation of a poly (A) tail. Nature structural & molecular biology 2019, 26, 1132–1140.

(4) Tesina, P.; Lessen, L. N.; Buschauer, R.; Cheng, J.; Wu, C. C.-C.; Berninghausen, O.; Buskirk, A. R.; Becker, T.; Beckmann, R.; Green, R. Molecular mechanism of translational stalling by inhibitory codon combinations and poly (A) tracts. The EMBO Journal 2020, 39, e103365.

(5) Seidelt, B.; Innis, C. A.; Wilson, D. N.; Gartmann, M.; Armache, J.-P.; Villa, E.; Trabuco, L. G.; Becker, T.; Mielke, T.; Schulten, K.; Steitz, T. A.; Beckmann, R. Structural Insight into Nascent Polypeptide Chain–Mediated Translational Stalling. Science 2009, 326, 1412–1415.

(6) Bischoff, L.; Berninghausen, O.; Beckmann, R. Molecular Basis for the Ribosome Functioning as an L-Tryptophan Sensor. Cell Reports 2014, 9, 469–475.

(7) Del Valle, A. H.; Seip, B.; Cervera-Marzal, I.; Sacheau, G.; Seefeldt, A. C.; Innis, C. A. Ornithine capture by a translating ribosome controls bacterial polyamine synthesis. Nature microbiology 2020, 5, 554–561.

(8) van der Stel, A.-X.; Gordon, E. R.; Sengupta, A.; Martinez, A. K.; Klepacki, D.; Perry, T. N.; del Valle, A. H.; Vazquez-Laslop, N.; Sachs, M. S.; Cruz-Vera, L. R.; et al. Structural basis for the tryptophan sensitivity of TnaC-mediated ribosome stalling. bioRxiv 2021,

(9) Chiba, S.; Lamsa, A.; Pogliano, K. A Ribosome–Nascent Chain Sensor of Membrane Protein Biogenesis in Bacillus Subtilis. The EMBO Journal 2009, 28, 3461–3475.

(10) Ishii, E.; Chiba, S.; Hashimoto, N.; Kojima, S.; Homma, M.; Ito, K.; Akiyama, Y.; Mori, H. Nascent Chain-Monitored Remodeling of the Sec Machinery for Salinity Adaptation of Marine Bacteria. Proceedings of the National Academy of Sciences 2015, 112, E5513–E5522.

(11) Kannan, K.; Kanabar, P.; Schryer, D.; Florin, T.; Oh, E.; Bahroos, N.; Tenson, T.; Weissman, J. S.; Mankin, A. S. The general mode of translation inhibition by macrolide antibiotics. Proceedings of the National Academy of Sciences 2014, 111, 15958–15963.

(12) Sohmen, D.; Chiba, S.; Shimokawa-Chiba, N.; Innis, C. A.; Berninghausen, O.; Beckmann, R.; Ito, K.; Wilson, D. N. Structure of the Bacillus Subtilis 70S Ribosome Reveals the Basis for Species-Specific Stalling. Nature Communications 2015, 6, 6941.

(13) Zhang, J.; Pan, X.; Yan, K.; Sun, S.; Gao, N.; Sui, S.-F. Mechanisms of Ribosome Stalling by SecM at Multiple Elongation Steps. eLife 2015, 4, e09684.

(14) Arenz, S.; Meydan, S.; Starosta, A. L.; Berninghausen, O.; Beckmann, R.; VázquezLaslop, N.; Wilson, D. N. Drug sensing by the ribosome induces translational arrest via active site perturbation. Molecular cell 2014, 56, 446–452.

(15) Arenz, S.; Bock, L. V.; Graf, M.; Innis, C. A.; Beckmann, R.; Grubmüller, H.; Vaiana, A. C.; Wilson, D. N. A Combined Cryo-EM and Molecular Dynamics Approach Reveals the Mechanism of ErmBL-Mediated Translation Arrest. Nature Communications 2016, 7, comms12026.

(16) Su, T.; Cheng, J.; Sohmen, D.; Hedman, R.; Berninghausen, O.; von Heijne, G.; Wilson, D. N.; Beckmann, R. The Force-Sensing Peptide VemP Employs Extreme Compaction and Secondary Structure Formation to Induce Ribosomal Stalling. eLife 2017, 6, e25642.

(17) Gumbart, J.; Schreiner, E.; Wilson, D. N.; Beckmann, R.; Schulten, K. Mechanisms of SecM-mediated stalling in the ribosome. Biophysical journal 2012, 103, 331–341.

(18) Bock, L. V.; Kolář, M. H.; Grubmüller, H. Molecular Simulations of the Ribosome and Associated Translation Factors. Current Opinion in Structural Biology 2018, 49, 27–35.

(19) Zimmer, M. H.; Niesen, M. J.; Miller, T. F. Force transduction creates long-ranged coupling in ribosomes stalled by arrest peptides. bioRxiv 2020,

(20) Miyazaki, R.; Akiyama, Y.; Mori, H. Fine interaction profiling of VemP and mechanisms responsible for its translocation-coupled arrest-cancelation. Elife 2020, 9, e62623.

(21) Mori, H.; Sakashita, S.; Ito, J.; Ishii, E.; Akiyama, Y. Identification and Characterization of a Translation Arrest Motif in VemP by Systematic Mutational Analysis. Journal of Biological Chemistry 2018, jbc.M117.816561.

(22) Nakatogawa, H.; Ito, K. The Ribosomal Exit Tunnel Functions as a Discriminating Gate. Cell 2002, 108, 629–636.

(23) Zhang, J.; Pan, X.; Yan, K.; Sun, S.; Gao, N.; Sui, S.-F. Mechanisms of ribosome stalling by SecM at multiple elongation steps. Elife 2015, 4, e09684.

(24) Lu, J.; Deutsch, C. Folding zones inside the ribosomal exit tunnel. Nature structural & molecular biology 2005, 12, 1123–1129.

(25) Ziv, G.; Haran, G.; Thirumalai, D. Ribosome exit tunnel can entropically stabilize α-helices. Proceedings of the National Academy of Sciences 2005, 102, 18956–18961.

(26) Sorin, E. J.; Pande, V. S. Nanotube confinement denatures protein helices. Journal of the American Chemical Society 2006, 128, 6316–6317.

(27) Fischer, N.; Neumann, P.; Konevega, A. L.; Bock, L. V.; Ficner, R.; Rodnina, M. V.; Stark, H. Structure of the E. Coli Ribosome-EF-Tu Complex at <3 A Resolution by Cs-Corrected Cryo-EM. Nature 2015, 520, 567–570.

(28) Maier, J. A.; Martinez, C.; Kasavajhala, K.; Wickstrom, L.; Hauser, K. E.; Simmerling, C. ff14SB: improving the accuracy of protein side chain and backbone parameters from ff99SB. Journal of chemical theory and computation 2015, 11, 3696–3713.

(29) Aduri, R.; Psciuk, B. T.; Saro, P.; Taniga, H.; Schlegel, H. B.; SantaLucia, J. AMBER Force Field Parameters for the Naturally Occurring Modified Nucleosides in RNA. Journal of Chemical Theory and Computation 2007, 3, 1464–1475.

(30) Berendsen, H. J. C.; Postma, J. P. M.; van Gunsteren, W. F.; Hermans, J. In Intermolecular Forces: Proceedings of the Fourteenth Jerusalem Symposium on Quantum Chemistry and Biochemistry Held in Jerusalem, Israel, April 13–16, 1981; Pullman, B., Ed., The Jerusalem Symposia on Quantum Chemistry and Biochemistry; Springer Netherlands: Dordrecht, 1981; pp 331–342.

(31) Joung, I. S.; Cheatham III, T. E. Determination of alkali and halide monovalent ion parameters for use in explicitly solvated biomolecular simulations. The journal of physical chemistry B 2008, 112, 9020–9041.

(32) Warias, M.; Grubmüller, H.; Bock, L. V. tRNA dissociation from EF-tu after GTP hydrolysis: primary steps and antibiotic inhibition. Biophysical journal 2020, 118, 151–161.

(33) Huang, J.; Rauscher, S.; Nawrocki, G.; Ran, T.; Feig, M.; de Groot, B. L.; Grubmüller, H.; MacKerell Jr, A. D. CHARMM36m: An Improved Force Field for Folded and Intrinsically Disordered Proteins. Nature Methods 2017, 14, 71–73.

(34) Jorgensen, W. L.; Chandrasekhar, J.; Madura, J. D.; Impey, R. W.; Klein, M. L. Comparison of Simple Potential Functions for Simulating Liquid Water. The Journal of Chemical Physics 1983, 79, 926–935.

(35) Bussi, G.; Donadio, D.; Parrinello, M. Canonical Sampling through Velocity Rescaling. The Journal of Chemical Physics 2007, 126, 014101.

(36) Berendsen, H. J. C.; Postma, J. P. M.; van Gunsteren, W. F.; DiNola, A.; Haak, J. R. Molecular Dynamics with Coupling to an External Bath. The Journal of Chemical Physics 1984, 81, 3684–3690.

(37) Parrinello, M.; Rahman, A. Polymorphic transitions in single crystals: A new molecular dynamics method. Journal of Applied physics 1981, 52, 7182–7190.

(38) Darden, T.; York, D.; Pedersen, L. Particle Mesh Ewald: An N·log(N) Method for Ewald Sums in Large Systems. The Journal of Chemical Physics 1993, 98, 10089– 10092.

(39) Hess, B. P-LINCS: A Parallel Linear Constraint Solver for Molecular Simulation. Journal of Chemical Theory and Computation 2008, 4, 116–122.

(40) Feenstra, K. A.; Hess, B.; Berendsen, H. J. Improving efficiency of large time-scale molecular dynamics simulations of hydrogen-rich systems. Journal of Computational Chemistry 1999, 20, 786–798.

(41) Abraham, M. J.; Murtola, T.; Schulz, R.; Páll, S.; Smith, J. C.; Hess, B.; Lindahl, E. GROMACS: High Performance Molecular Simulations through Multi-Level Parallelism from Laptops to Supercomputers. SoftwareX 2015, 1-2, 19–25.

(42) Kabsch, W.; Sander, C. Dictionary of Protein Secondary Structure: Pattern Recognition of Hydrogen-Bonded and Geometrical Features. Biopolymers 1983, 22, 2577– 2637.

(43) Nagy, G.; Igaev, M.; Jones, N. C.; Hoffmann, S. V.; Grubmüller, H. SESCA: predicting circular dichroism spectra from protein molecular structures. Journal of chemical theory and computation 2019, 15, 5087–5102.

(44) Nagy, G.; Grubmüller, H. Implementation of a Bayesian Secondary Structure Estimation Method for the SESCA Circular Dichroism Analysis Package.

(45) Cuff, J. A.; Barton, G. J. Application of Multiple Sequence Alignment Profiles to Improve Protein Secondary Structure Prediction. Proteins: Structure, Function, and Bioinformatics 2000, 40, 502–511.

(46) Drozdetskiy, A.; Cole, C.; Procter, J.; Barton, G. J. JPred4: A Protein Secondary Structure Prediction Server. Nucleic Acids Research 2015, 43, W389–W394.

(47) Combet, C.; Blanchet, C.; Geourjon, C.; Deléage, G. NPS@: Network Protein Sequence Analysis. Trends in Biochemical Sciences 2000, 25, 147–150.

(48) Petrone, P. M.; Snow, C. D.; Lucent, D.; Pande, V. S. Side-chain recognition and gating in the ribosome exit tunnel. Proceedings of the National Academy of Sciences 2008, 105, 16549–16554.

(49) Lucent, D.; Snow, C. D.; Aitken, C. E.; Pande, V. S. Non-bulk-like solvent behavior in the ribosome exit tunnel. PLoS Comput Biol 2010, 6, e1000963.

(50) Makarov, G.; Golovin, A.; Sumbatyan, N.; Bogdanov, A. Molecular dynamics investigation of a mechanism of allosteric signal transmission in ribosomes. Biochemistry (Moscow) 2015, 80, 1047–1056.

(51) Tsai, C.-J.; Del Sol, A.; Nussinov, R. Protein allostery, signal transmission and dynamics: a classification scheme of allosteric mechanisms. Molecular Biosystems 2009, 5, 207–216.

(52) Samatova, E.; Daberger, J.; Liutkute, M.; Rodnina, M. V. Translational control by ribosome pausing in bacteria: how a non-uniform pace of translation affects protein production and folding. Frontiers in Microbiology 2020, 11.

